# gJLS2: An R package for generalized joint location and scale analysis in X-inclusive genome-wide association studies

**DOI:** 10.1101/2021.10.11.463951

**Authors:** Wei Q. Deng, Lei Sun

## Abstract

A joint analysis of location and scale can be a powerful tool in genome-wide association studies to uncover previously overlooked markers that influence a quantitative trait through both mean and variance, as well as to prioritize candidates for gene-environment interactions. This approach has recently been generalized to handle related samples, dosage data, and the analytically challenging X-chromosome. We disseminate the latest advances in methodology through a user-friendly R software package with added functionalities to support genome-wide analysis on individual-level or summary-level data. The implemented R package can be called from PLINK or directly in a scripting environment, to enable a streamlined genome-wide analysis for biobank-scale data. Application results on individual-level and summary-level data highlight the advantage of the joint test to discover more genome-wide signals as compared to a location or scale test alone. We hope the availability of gJLS2 software package will encourage more scale and/or joint analyses in large-scale datasets, and promote the standardized reporting of their *p*-values to be shared with the scientific community.

## 1 Introduction

Genetic association studies examine the relationship between the genotypes of a single-nucleotide polymorphism (SNP; denoted by *G*) and a quantitative phenotype (denoted by *Y*), by testing the mean differences in *Y* according to the genotypes, otherwise known as a location test. More recently, several reports have investigated association with phenotypic variance of complex quantitative traits (Pare, Cook, Ridker, & Chasman, 2010; Shungin et al., 2017; Yang et al., 2012), by testing the variance differences in *Y* across the genotype groups of a SNP, or known as a scale test, in hopes of finding biologically meaningful markers. One possible explanation of a significant scale effect is the presence of gene–gene (GxG) or gene–environment (GxE) interactions; both referred to as GxE hereinafter. Unlike a direct test of GxE interaction, a scale test can be used to indirectly infer GxE without knowledge of the interacting covariates, thus alleviates the multiple hypothesis burden of testing all possible pairwise interactions, and the assumption that all interacting environmental variables could be (accurately) measured.

A more powerful approach to prioritize biologically relevant candidates is to jointly evaluate location and scale effects (Aschard, Hancock, London, & Kraft, 2010; Cao et al., 2014; Soave et al., 2015). As compared to other existing single-marker based joint tests, a joint location-scale (JLS) association test (Soave et al., 2015) is easy to implement on individual-level data or in a meta-analysis, requiring only the location and scale *p*-values for each SNP, marking the first generation of the JLS tool (https://github.com/dsoave/JLS). Since methods for genome-wide association studies (GWAS) of location are well-established, the main focus was on improving scale tests tailored for genetic data. Soave et al. (2017) generalized the well-known Levene’s test to handle complex data structures often observed in genetic studies, such as correlated samples and dosage data, leading to the next update, namely, generalized JLS (gJLS; https://github.com/dsoave/gJLS). Following the X-inclusive trend to genome-wide analyses, robust and powerful location (Chen et al, 2021) and scale tests (Deng et al., 2019) tailored for X-chromosome are now available.

In this paper, we describe a generalized joint location and scale analysis tool (gJLS2) as an update to the JLS and gJLS methodology for autosomes, with added functionalities that 1) support X-chromosome mean and variance association analyses, 2) handle imputed data as genotypic probabilities or in dosage format, 3) allow the incorporation of summary statistics for location and/or scale tests, 4) implement a flexible framework that can accommodate additional covariates in both the location and scale association models, and 5) improve the computational time required for large-scale genetic data such as the UK biobank (Bycroft et al., 2018). We hope the availability of this unifying software package will encourage more X-inclusive, genome-wide, gJLS2 association analysis for complex continuous traits, particularly for those believed to be enriched for genetic interactions.

## 2 Methods

The software can be installed in an R environment from CRAN:

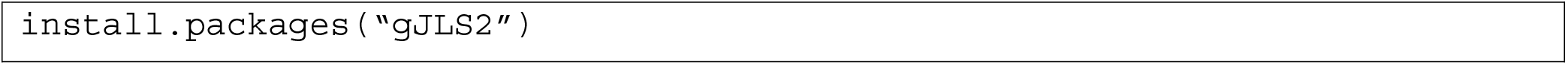

or github for the most recent version:

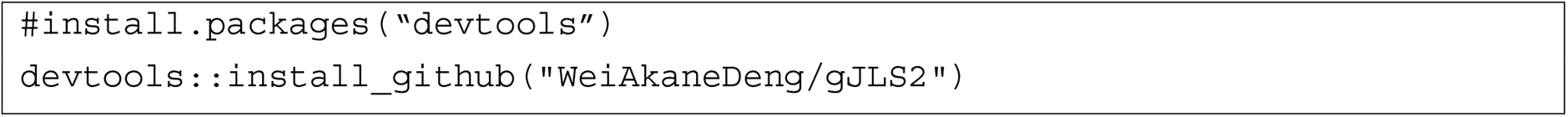

via the “devtools” package (Wickham et al., 2018).

### 2.1 Data preparation

The gJLS2 software package requires the minimal inputs of a quantitative trait and genotype data, which can be discrete genotype counts, continuous dosage genotype values or imputed genotype probabilities. The genotype data can be supplied in PLINK format via the R plug-in option using PLINK 1.9 (Chang et al., 2015) or any other format that can be read in R with packages *“BGData”* and *“BEDMatrix”* (Grueneberg and Campos, 2019). The phenotype and covariate should be supplied in the same file, and if sex is required for X-chromosome analysis, then it needs to be the first column after individual IDs.

For smaller analyses or testing purpose, it is possible to use the package in R GUI directly. However, for genome-wide analyses on larger datasets, we recommend either Rscript commands or as an R-plugin within PLINK combined with a high-performing computing cluster environment. We will demonstrate all three approaches using the example datasets provided.

### 2.1 Example datasets

The package included two example datasets: one comprises of simulated phenotype, covariate data and real X-chromosome genotypes from the 1000 Genomes Project (The 1000 Genomes Project Consortium, 2015), denoted by “chrXdat”; and another is based on summary statistics from the Genetic Investigation of ANthropometric Traits (GIANT) consortium for body mass index (BMI) and human standing height (Yang et al., 2012; Yengo et al., 2018).

The dataset “chrXdat” consists of 5 hand-picked SNPs rs5983012 (A/G), rs986810 (C/T), rs180495 (G/A), rs5911042 (T/C), and rs4119090 (G/A) that are outside of the pseudo-autosomal region, to cover observed minor allele frequency (MAF; calculated in females and rounded to the nearest digit) of 0.1, 0.2, 0.3, 0.4 and 0.45, respectively. See Supplementary Material Section 2 for more details on the simulated dataset. The summary statistics data comprised of association *p*-values of SNPs with the *mean* of BMI and height, under the column name “gL” for location, and those with the *variance*, under the column name “gS”. The example data only included chromosome 16 SNPs to keep the size of example datasets manageable.

### 2.2 Location association

For autosomal SNPs, a linear regression fitted using the ordinary least square (OLS) is flexible to accommodate additional covariates and the default option. To account for related samples, the generalized least square (GLS) method is used by specifying a covariance structure for error terms in smaller samples. Users can either provide the covariance matrix or specify a structure for the covariance matrix according to pre-defined subgroups. However, for large population studies (*n* > 5,000), it is recommended to run the location association analysis using the state-of-art linear mixed models, such as LMM-BOLT (Loh *et al*., 2015) or SAIGE (Zhou *et al*., 2018) and supply the results as location summary statistics for the joint analysis.

The novel contribution of gJLS2 is the addition of our recommended X-chromosome location association strategy (Chen *et al*., 2021), which has good type I error control, is robust to sex confounding, arbitrary baseline allele choice, uncertainty of X-Chromosome Inactivation (XCI), and skewed XCI. The X-chromosome association requires additionally the sex information and the default location test *p*-value is obtained by testing the null hypothesis **H**_o_: β_G_ = β_GS_ = *β_D_* = 0 under the linear model:

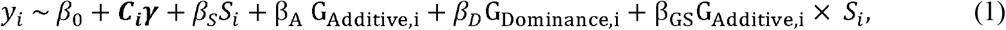

where *G_A_* is the additively coded genotype variable, *G_D_* is an indicator variable for the female heterozygotes, the sex variable S is coded with males as the baseline taking value 0 and females taking value 1, and *C* is a vector of covariates to be adjusted for. The regression coefficients *β*_0_, **γ**, *β_S_* are for the non-genetic covariates in the model and β_G_, β_GS_, *β_D_* denote the regression coefficients of interest, for the additive, GxSex, and dominance effects, respectively. The function “locReg” can be called with the X-chromosome option and returns the default 3-degree of freedom (df) test *p*-value:

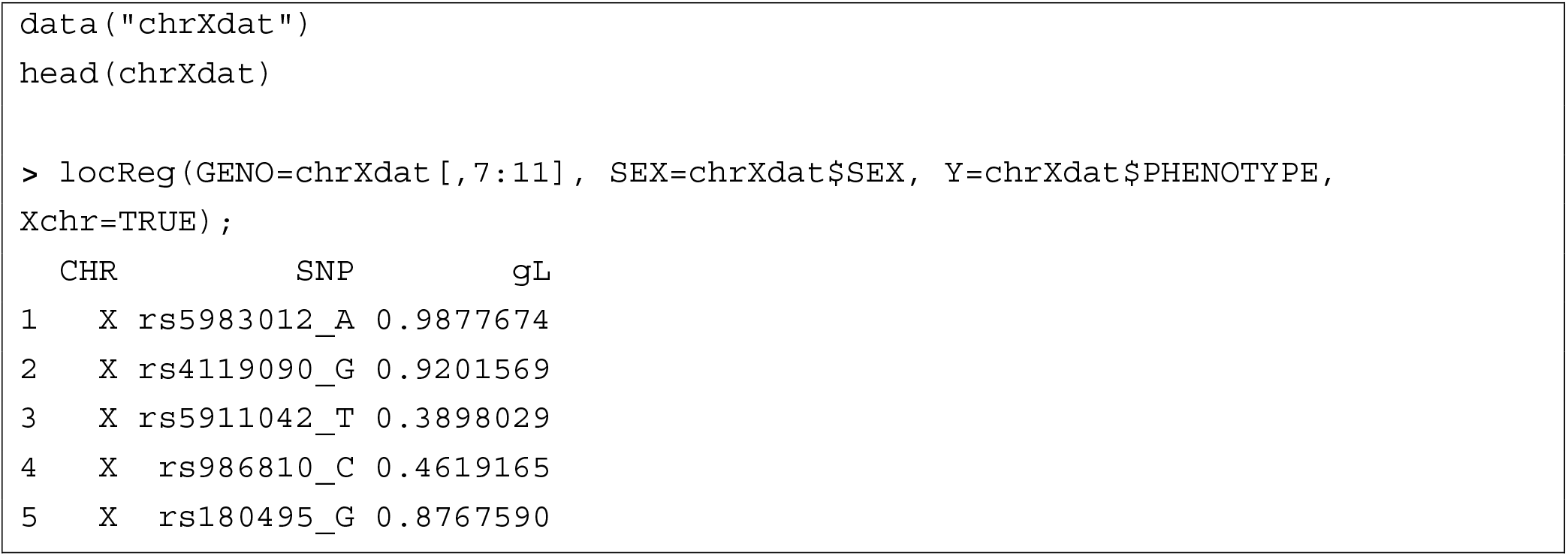

Although the resulting 3-df test is robust to the choice of baseline allele and status of XCI, but a recent report (Pazokitoroudi *et al*., 2021) has found “limited contribution of dominance hereditability to complex trait variation” and the recommended 3-df test can only be applied to discrete genotypes. Thus, an alternative 2-df test without the dominance term is recommended and the default option for imputed SNPs.

### 2.3 Scale association

The scale component builds on two recent works that generalized to dosage genotype, related samples (Soave et al., 2017), and the X-chromosome (Deng *et al*., 2019), both are extensions of the generalized Levene’s test via a regression framework. Besides a more flexible characterization of sample dependence structure, the generalization also allows analysis of imputed genotype data, which would otherwise be challenging with a group-based variance heterogeneity test, e.g. the Levene’s test or the Brown-Forsythe test.

The variance test *p*-value is obtained using a two-stage generalized Levene’s test assuming additive variance effects. Briefly, the residuals *d* was computed under Equation (1) using the Least Absolute Deviation (LAD) algorithm, which gives the residuals in terms of each observation’s distance with respect to the median (rather than the mean as in OLS) and is the default option. The absolute residuals are then fitted under the 3-df recommended model for discrete genotype:

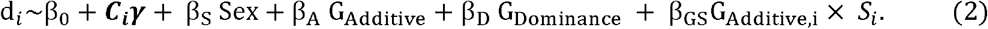

The following examples show results from LAD and OLS algorithms are very similar when the simulated phenotype is roughly symmetric:

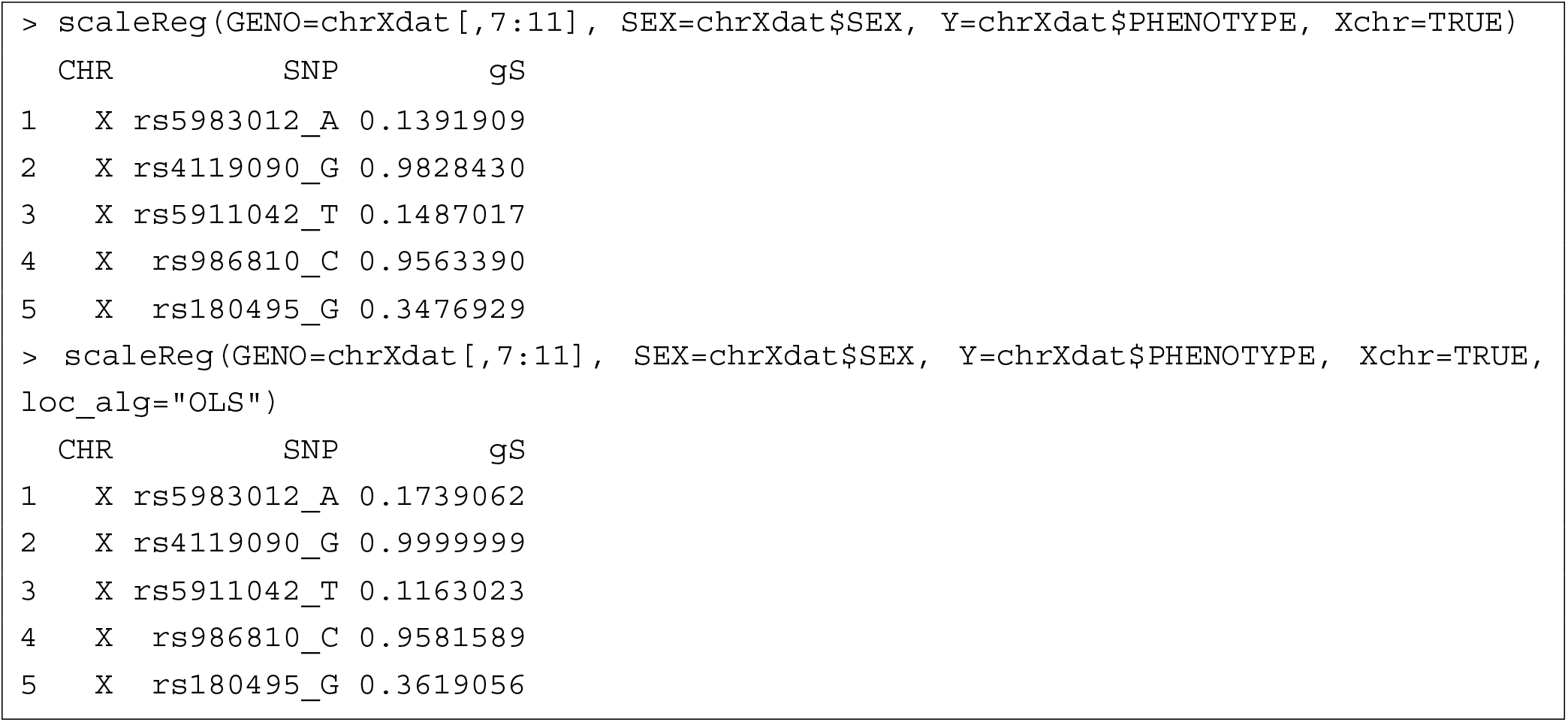

The imputed data are analyzed either by computing the dosage value and used in place of *G* additively (2 df) without the dominance term, or by replacing the genotype indicators for each observation, with the corresponding group probabilities (3 df). Similar to the location association, sample relatedness is dealt with using generalized least squares (GLS) for autosomal markers at the second stage of linear regression via the correlation matrix. Additional models, such as a sex-stratified variance test, can also be specified for X-chromosome. In this case, the scale test result is given by the Fisher’s method that combines female and male-specific variance test results.

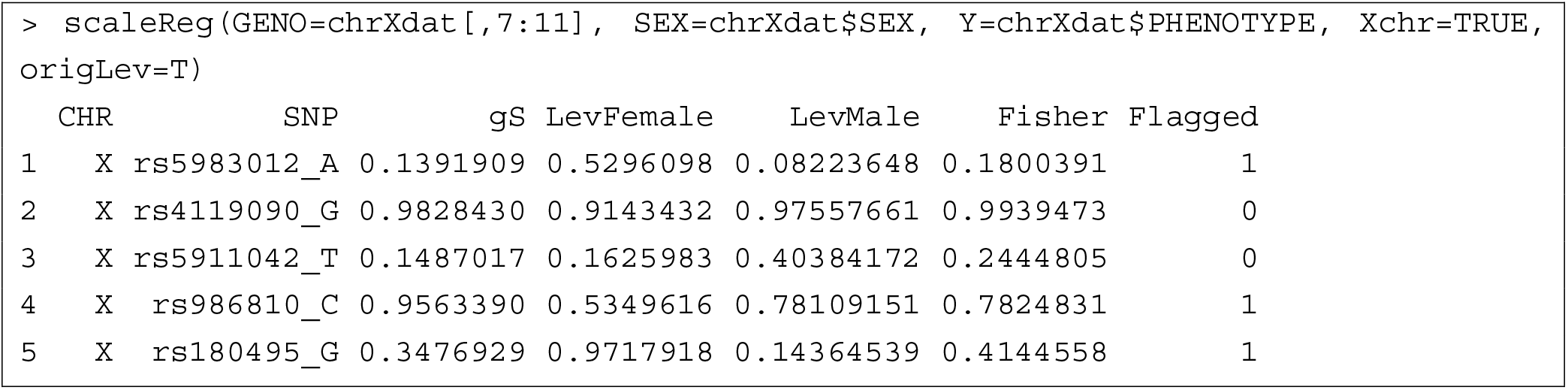

Note that a “Flagged” column is appended for these results, indicating the minimum genotype count in either females/males is less than 30 (indicated by a value of 1) or not (indicated by 0). This is based on the quality control that SNPs with a minimum count below 30 should be removed to avoid inflated type I errors (Deng et al., 2019; Soave et al., 2015).

### 2.4 Joint location-and-scale analysis

The joint analyses can be performed automatically as part of the *gJLS2* pipeline after running location and scale tests, where the default option applies a quantile transformation to the quantitative trait such that the location and scale test can be combined without inflating the type I error rates.

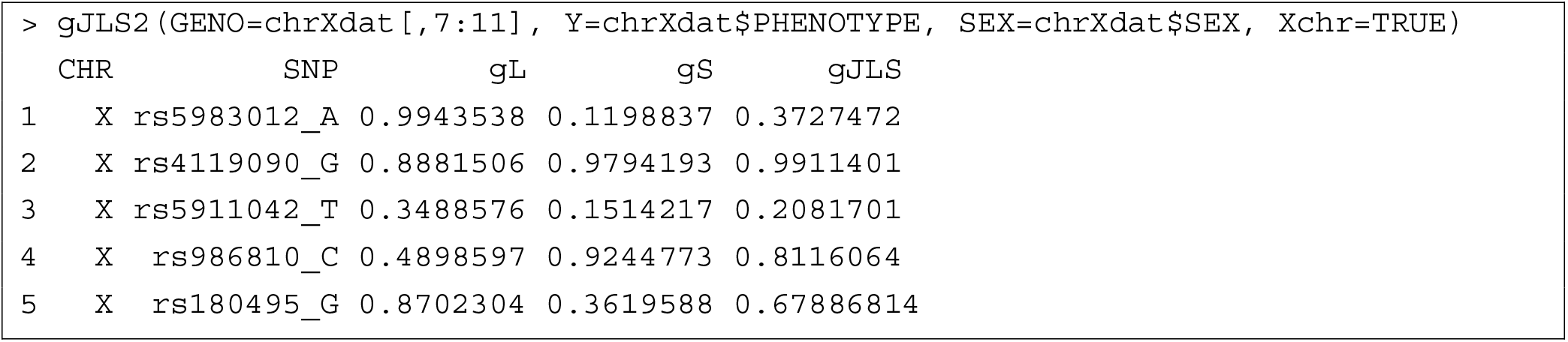

For larger datasets, it is more convenient to run the joint analyses using the PLINK R plug-in following the a typical GWAS pipeline. The R plug-in relies on the *Rserve* package to establish communication between R and PLINK 1.9. The following script demonstrates the joint analyses for X-chromosome SNPs that included additional covariates.

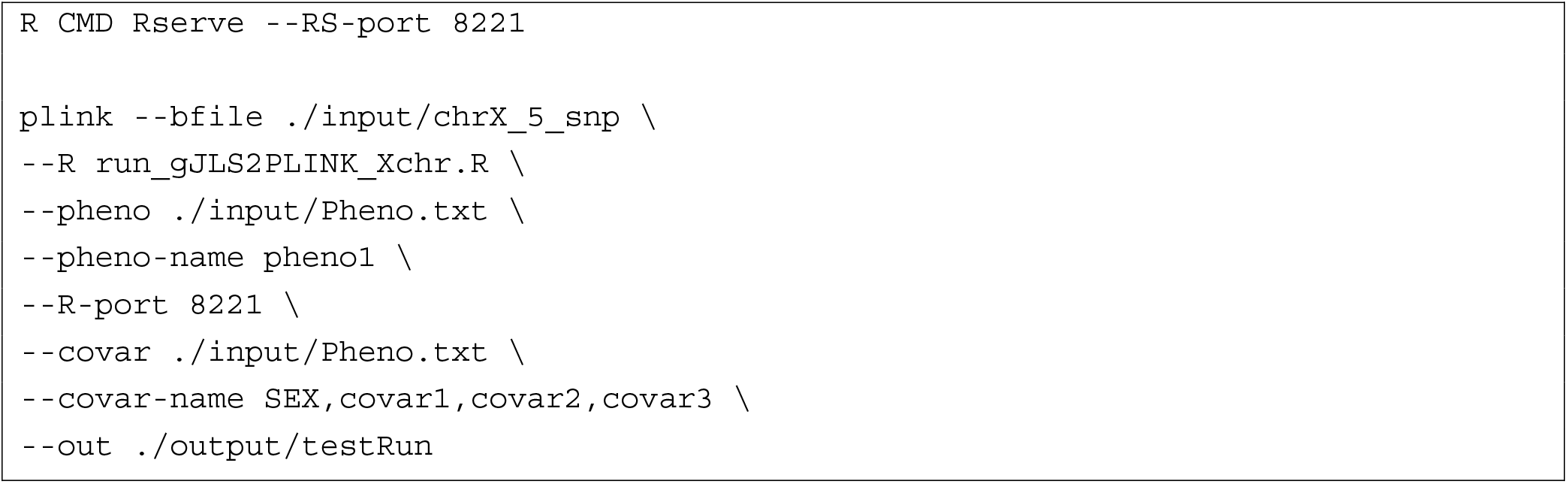

Another option is to use the Rscript provided that allows additional arguments to increase the number of cores used (--nThreads) and to change how frequently the results are written to the output (--write).

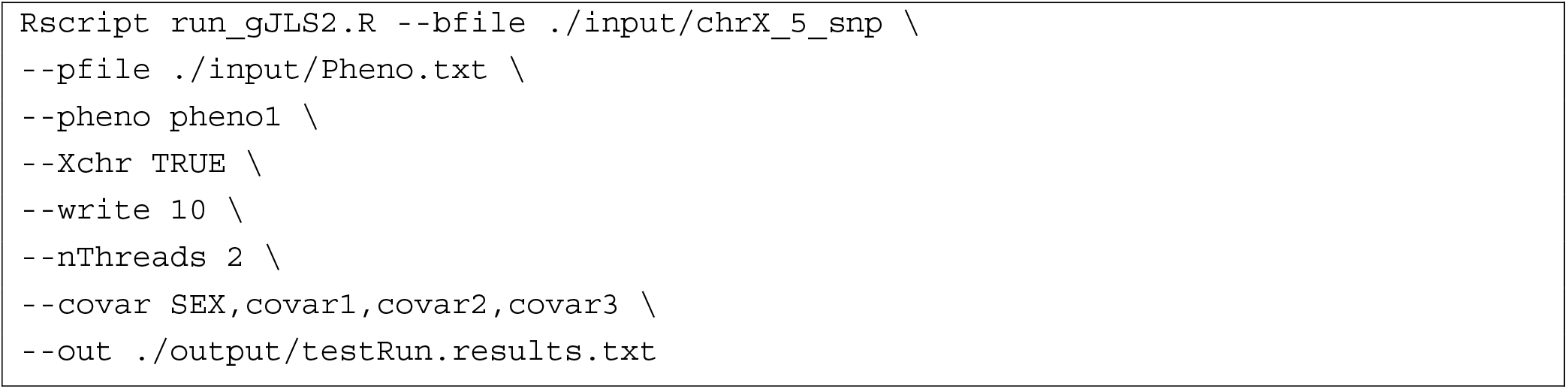

Alternatively, summary statistics, i.e. *p*-values from location and scale tests are allowed and can be combined via Fisher’s method to give the corresponding test statistic for the joint analysis:

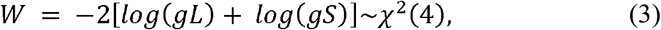

where *gL* denotes the location *p*-value and *gS* the scale *p*-value.

An important assumption underlying this simple method to combine evidence is the normality of the quantitative trait, which leads to *gL* and *gS* being independent under the null of no association. Though empirically the analyses remain valid for approximated normal distributions (Soave et al., 2017), we strongly recommend the user to follow the default option and quantile transform the quantitative trait for location and scale association. We expect the independence between *gL* and *gS* to hold for X-chromosomal SNPs under appropriate location and scale models that accounted for the confounding effect of sex, XCI uncertainty and skewed XCI (Chen *et al*., 2021). The simulation study in Section 2 of the Supplementary Material was conducted to help readers gauge the effect of non-normality on the gJLS *p*-values for X-chromosome markers.

## 3 Application of gJLS2 to real datasets

The *gJLS2* software supports the joint analysis on both individual-level data as well as summary statistics. To highlight the functions in our package, we present 1) an X-chromosome gJLS analysis on UK Biobank (UKB) data (Bycroft et al., 2018) on four complex traits previously studied in Deng et al. (2019); 2) a chromosomewide gJLS analysis on summary statistics of location and scale from the GIANT consortium for BMI and height.

### 3.1 Application to UK Biobank data

The sex-stratified means and variances for these quantitative traits are summarized graphically in **Supplementary Figures 1-4**. A quick visual inspection suggests that quantile normalization should be applied, which is also the default options in *gJLS2*. We restricted analyses to white British samples (n = 276,694), and for BMI, we further excluded those with diagnosed type 2 diabetes (n = 262,837). We included only bi-allelic SNPs and filtered based on MAF < 0.01, HWE *p*-value < 1E-5. A further check on sex-stratified MAFs (**Suppl. Figure 5**) confirmed that the remaining 15,179 X-chromosomal SNPs are of good quality.

The genotype data can be supplied in PLINK binary format via the R plug-in option using PLINK 1.9 (Chang *et al*., 2015) or any other format that can be read in R with packages *“BGData”* and *“BEDMatrix”* (Grueneberg and Campos, 2019). The phenotype and covariate should ideally be supplied in the same file, and if not, should have a common column to match samples. For genome-wide analyses on larger datasets, we recommend the use of a high-performing computing cluster and taking advantage of multiple cores whenever available.

For the location and scale regression, we included age, genetic sex, and the first 10 genetic principal components. Since there is no additional arguments needed, such as dosage or related samples, the analysis can be done using either a PLINK R-plugin running on PLINK 1.9:

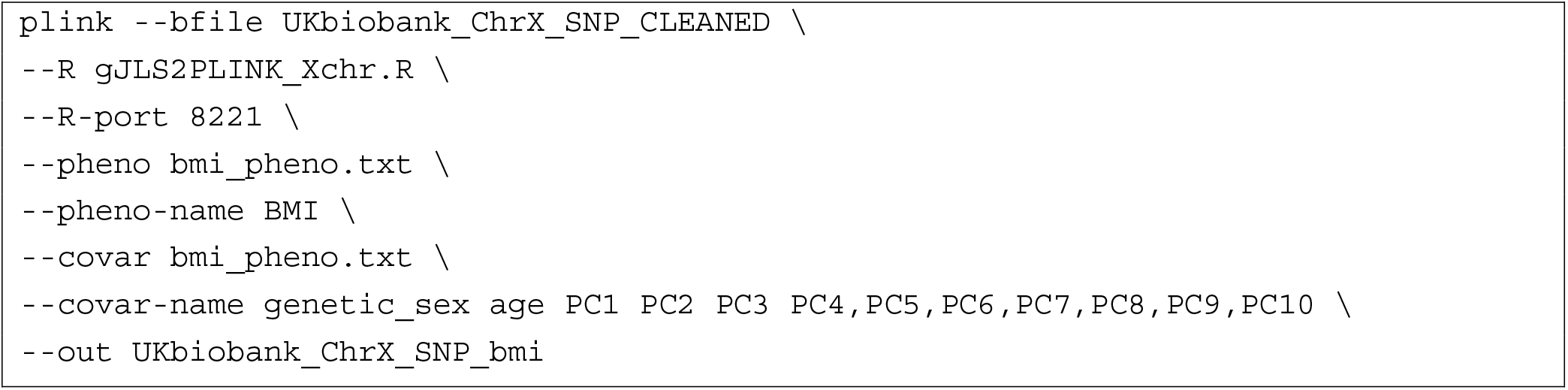

or the Rscript option:

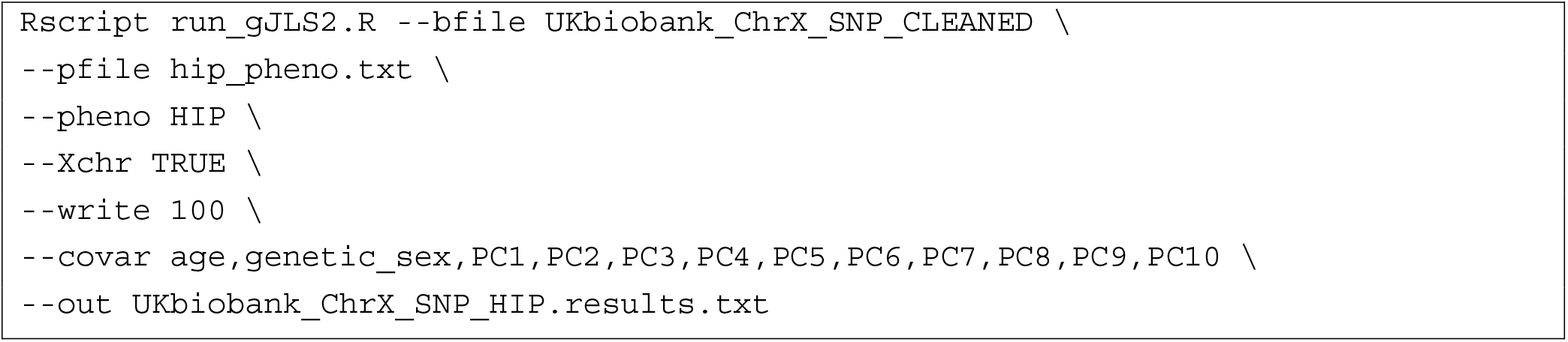

The base scripts for PLINK (`gJLS2PLINK_Xchr.R`) and Rscript (`run_gJLS2.R`) are provided in the inst/extdata folder of the R package along with the input files. It is worth noting the main advantages of the Rscript option beyond its flexibility: 1) the gJLS2 R package supports multi-core computing via the *parallel* base package and the argument “nThreads” can be used to specify the number of cores; 2) another useful feature is the “write” option that specifies the chunk size for the results to be written while the analysis is running and thus minimizes loss in case an interruption occurred.

Since the PLINK plug-in option only supports single-thread computation, we fixed the number of cores to be 1. There is no drastic difference between the two options, both took ~20 hours with 20G allocated memory (computing node specs 2xIntel E5-2683 v4 Broadwell @ 2.1GHz), to complete the analysis for 276,694 unrelated European samples and 15,179 X-chromosome variants per trait. The gL, gS, and gJLS2 *p*-values are presented using a Manhattan plot, quantile-quantile plot, and a histogram (**Suppl Figures 6-9**) for each of the complex traits. We also tabulated a list of SNPs that did not pass the genome-wide significance threshold of 5E-8 for gL, but did pass for gJLS in **Table *1***, demonstrating the benefit of gJLS for additional genome-wide discoveries.

**Table 1.**
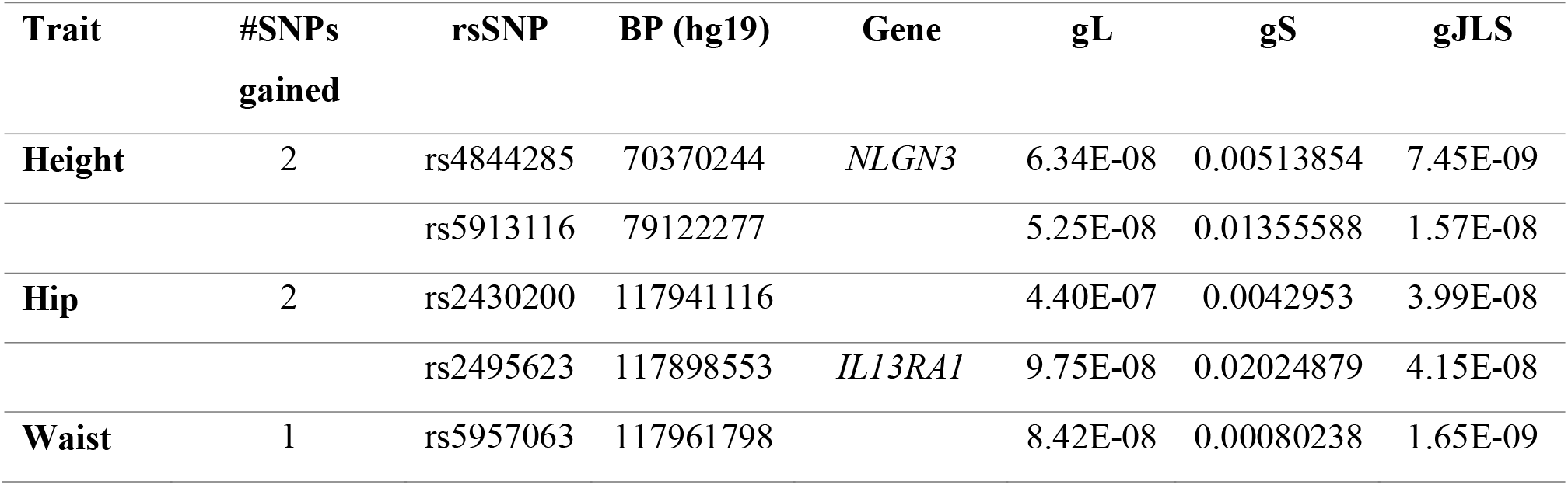
Number of X-chromosomal SNPs gained at genome-wide significance by the joint location-scale *p*-values for four complex traits in UKB.

### 4.2 Application to summary statistics

The Rscript is more flexible than the PLINK plug-in solution as it can also handle analysis of summary statistics. The input file should contain at least three columns with headers “SNP”, “gL”, and “gS”, while the output file has an additional column “gJLS”. We re-formatted the subset of chromosome 16 summary statistics of location and scale obtained from the GIANT consortium for body mass index (BMI) and height as inputs.

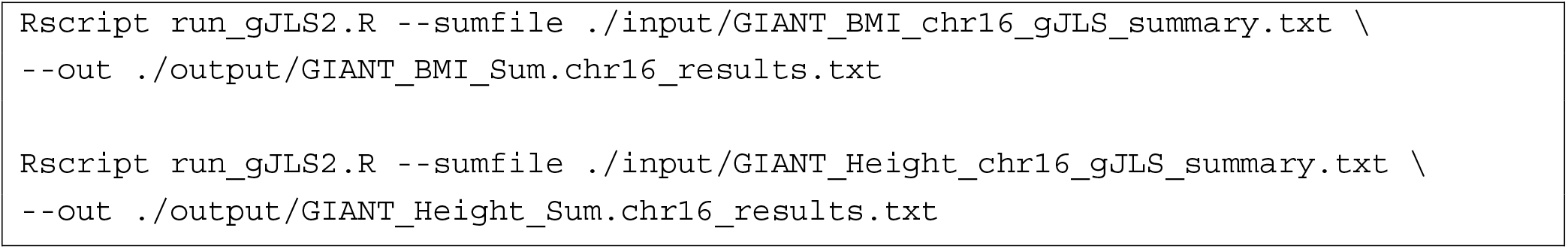

The gL, gS, and gJLS2 *p*-values are presented using a Manhattan plot, quantile-quantile plot, and a histogram (**Suppl Figures 10-11**). For chromosome 16, we gained 2 SNPs in *DNAH3* gene that passed the genome-wide significance using gJLS for BMI and an additional 23 SNPs for height (**Supplementary Table 1**).

## 4 Concluding remarks

The gJLS2 software package is a versatile tool for genome-wide discovery that is X-chromosome inclusive. As compared to previous versions, namely, JLS and gJLS, it has improved remarkably in terms of methodology, flexibility, computational improvements and most importantly, usability for large-scale data. Meanwhile, we expect many new features to be added with ongoing work. Naturally, the analysis of rare variant is a possible future direction for scale association test under the regression framework of a generalized Levene’s test. With the available summary statistics, a systematic approach to prioritize SNPs that takes into account the joint-location, scale effects, and functional annotation is needed to produce relevant candidates for gene-environment interaction. Finally, apart from the improved signal detection, the unbiased estimation of location effects for X-chromosome and scale effects in general remain open questions, but are expected to yield improved performance of polygenic prediction.

## Data Availability Statement

The gJLS2 package and the example datasets are available from https://github.com/WeiAkaneDeng/gJLS2. The simulations in this paper were based on X-chromosome genotypes from the 1000 Genomes Project (http://www.1000genomes.org), which can be freely accessed from the online data portal. The genotype and phenotype data used to demonstrate the X-chromosome analysis are available from UK Biobank (https://www.ukbiobank.ac.uk/) release in March 2018 and under project identification number 64875. Data access can be requested with information provided here: http://www.ukbiobank.ac.uk/using-the-resource/. The genome-wide summary statistics are made available from the GIANT data portal (https://portals.broadinstitute.org/collaboration/giant/index.php/GIANT_consortium_data_files).

## Acknowledgements

The authors would like to acknowledge Jiafen Gong, Sangook Kim, and Boxi Lin for testing and providing feedbacks on the initial versions of the gJLS2 software. The authors thank David Soave, the original creator of JLS and gJLS, for a critical reading of the paper.

---

**Suppl Fig. 1. Distribution of Body Mass Index (BMI) in 276,694 unrelated, white British samples from UKB.** The histograms (top figures) and the quantile-quantile plots (bottom figures) are presented for both the original scale and the inverse-normal transformed scale.

**Suppl Fig. 2. Distribution of height in 276,694 unrelated, white British samples from UKB.** The histograms (top figures) and the quantile-quantile plots (bottom figures) are presented for both the original scale and the inverse-normal transformed scale.

**Suppl Fig. 3. Distribution of hip circumference in 276,694 unrelated, white British samples from UKB.** The histograms (top figures) and the quantile-quantile plots (bottom figures) are presented for both the original scale and the inverse-normal transformed scale.

**Suppl Fig. 4. Distribution of waist circumference in 276,694 unrelated, white British samples from UKB.** The histograms (top figures) and the quantile-quantile plots (bottom figures) are presented for both the original scale and the inverse-normal transformed scale.

**Suppl Fig. 5. Comparison of Allele Frequency in Males and Females of the 15,179 X-chromosome markers from UK Biobank.**

**Suppl Fig. 6. XCHR-wide location, scale and gJLS test results for body mass index (BMI) using UK Biobank data.**

**Suppl Fig. 7. XCHR-wide location, scale and gJLS test results for height using UK Biobank data.**

**Suppl Fig. 8. XCHR-wide location, scale and gJLS test results for hip circumference using UK Biobank data.**

**Suppl Fig. 9. XCHR-wide location, scale and gJLS test results for waist circumference using UK Biobank data.**

**Suppl Fig. 10. Chromosome 16 location, scale and gJLS test results for BMI using summary statistics.**

**Suppl Fig. 11. Chromosome 16 location, scale and gJLS test results for height using summary statistics.**

## References

The 1000 Genomes Project Consortium. (2015). A global reference for human genetic variation. Nature. 526, 68–74. https://doi.org/10.1038/nature15393

Bycroft, C., Freeman, C., Petkova, D. et al. (2018). The UK Biobank resource with deep phenotyping and genomic data. Nature. 562, 203–209 https://doi.org/10.1038/s41586-018-0579-z

Christopher C Chang, Carson C Chow, Laurent CAM Tellier, Shashaank Vattikuti, Shaun M Purcell, James J Lee, (2015). Second-generation PLINK: rising to the challenge of larger and richer datasets, GigaScience, Volume 4, Issue 1, s13742-015-0047-8, https://doi.org/10.1186/s13742-015-0047-8.

Chen, B., Craiu, R. V., Strug, L. J., & Sun, L. (2021). The X factor: A robust and powerful approach to X-chromosome-inclusive whole-genome association studies. Genetic epidemiology, 10.1002/gepi.22422. Advance online publication. https://doi.org/10.1002/gepi.22422

Deng, W. Q., Mao, S., Kalnapenkis, A., Esko, T., Mägi, R., Paré, G., & Sun, L. (2019). Analytical strategies to include the X-chromosome in variance heterogeneity analyses: Evidence for trait-specific polygenic variance structure. Genetic epidemiology, 43(7), 815–830. https://doi.org/10.1002/gepi.22247.

Grueneberg A, de los Campos G (2019). “BGData - A Suite of R Packages for Genomic Analysis with Big Data.” G3: Genes, Genomes, Genetics, 9(5), 1377–1383. https://doi.org/10.1534/g3.119.400018.

Loh, PR., Tucker, G., Bulik-Sullivan, B. et al. (2015). Efficient Bayesian mixed-model analysis increases association power in large cohorts. Nature Genetics. 47, 284–290. https://doi.org/10.1038/ng.3190

Pazokitoroudi A, Chiu AM, Burch KS, Pasaniuc B, Sankararaman S. (2021) Quantifying the contribution of dominance deviation effects to complex trait variation in biobank-scale data. American Journal of Human Genetics. 108(5):799–808. https://doi.org/10.1016/j.ajhg.2021.03.018.

Roslin, N. M., Weili, L., Paterson, A. D. & Strug, L. J. (2016). Quality control analysis of the 1000 Genomes Project Omni2.5 genotypes. BioRxiv, https://doi.org/10.1101/078600

Soave, D., Corvol, H., Panjwani, N., Gong, J., Li, W., Boëlle, P. Y., Durie, P. R., Paterson, A. D., Rommens, J. M., Strug, L. J., & Sun, L. (2015). A Joint Location-Scale Test Improves Power to Detect Associated SNPs, Gene Sets, and Pathways. American journal of human genetics. 97(1), 125–138. https://doi.org/10.1016/j.ajhg.2015.05.015.

Soave D, Sun L. (2017). A generalized Levene’s scale test for variance heterogeneity in the presence of sample correlation and group uncertainty. Biometrics. 73(3):960–971. https://doi.org/10.1111/biom.12651.

Wickham, H., J. Hester, and W. Chang, (2018) devtools: Tools to Make Developing R Packages Easier. https://CRAN.R-project.org/package=devtools.

Yang J, Loos RJ, Powell JE, Medland SE, Speliotes EK, Chasman DI, Rose LM, Thorleifsson G, Steinthorsdottir V, Mägi R, et al. (2012). FTO genotype is associated with phenotypic variability of body mass index. Nature 490:267–272.

Yengo L, Sidorenko J, Kemper KE, Zheng Z, Wood AR, Weedon MN, Frayling TM, Hirschhorn J, Yang J, Visscher PM; GIANT Consortium. (2018) Meta-analysis of genome-wide association studies for height and body mass index in ~700000 individuals of European ancestry. Hum Mol Genet. 27(20):3641–3649. doi: 10.1093/hmg/ddy271. PMID: 30124842; PMCID:PMC6488973.

Zhou, W., Nielsen, J.B., Fritsche, L.G. et al. (2018). Efficiently controlling for case-control imbalance and sample relatedness in large-scale genetic association studies. Nature Genetics 50, 1335–1341. https://doi.org/10.1038/s41588-018-0184-y.

