## Supplementary Materials for "gJLS2: An R package for generalized joint location and scale analysis in X-inclusive genome-wide association studies"

### 1 Background Methods

Consider a quantitative trait *Y*, assumed to be (approximately) normally distributed or had been quantile transformed to resemble a normal distribution. Without loss of generality, the following linear model is assumed to reflect the ‘*true’* association relationship between *Y* and the SNP genotype variable *G*,

$Y = \beta_{0}+\beta_{G}G+\beta_{E}E+\beta_{GE}GE+\varepsilon$*,*	(1)

where *G* is coded additively with respect to the number of minor alleles taking discrete values 0, 1 and 2; *E* is a continuous covariate following the classical *G*-*E* independence assumption (Lindstrom et al., 2009); and the error term *ε* ∼ *N*(0, 1) is independent of *G* and *E*. Without observing *E*, the associated between *Y* and *G* is usually assessed via the *working* model of *Y* ~ *G.* Under the above assumptions, we observe that both the mean and variance of *Y* are expected to vary in the presence of GxE interactions:

$\text{E}\left( Y | G = g \right)=[\beta_{g}+\beta_{GE}\text{E}(E)]g$, (2)

$\text{Var}\left( Y | G = g \right)=\left( \beta_{E}+\beta_{GE}g \right)^{2}+ 1.$ (3)

The independence of the location and scale tests under the null hypothesis of no association has been previously established for autosomal SNPs assuming the error is normally distributed. As a result, the Fisher’s method can be used to combine the two independent *p*-values without concerns of inflated type I errors.

### 2 Simulated Dataset for an X-chromosome-wide gJLS Analysis

**Data simulation**

Instead of simulating the genotypes of X-chromosome (Xchr) SNPs, we had taken the Xchr genotypes from the 1000 Genomes Project while restricting to the European subset (*n* = 471 with non-ambiguous sex information). The cleaned 1000 Genomes Project genotype (Illumina Omni2.5M array) in PLINK format was obtained from <http://www.tcag.ca/tools/1000genomes.html> and a detailed report on the quality control procedure implemented can be found in Roslin et al., (2016). We hand-picked SNPs rs5983012 (A/G), rs986810 (C/T), rs180495 (G/A), rs5911042 (T/C), and rs4119090 (G/A) that are outside of the pseudo-autosomal region, to cover observed minor allele frequency (MAF; calculated in females and rounded to the nearest digit) of 0.1, 0.2, 0.3, 0.4 and 0.45, respectively.

Following the true model, we simulated the quantitative trait under the null hypothesis of no location nor scale effects:

$Y = \beta_{0}+\beta_{S}S+\beta_{E}E+\beta_{ES}ES+\varepsilon$, (1)

where we set parameter $\beta_{S}$ to be either 0, 0.5, -0.5 to mimic different sex-stratified *mean* conditions and $\beta_{ES}$ to be either 0, 0.3, -0.3 to mimic different sex-stratified *variance* conditions. The sex variable $S$ is coded with males as the baseline taking value 0 and females taking value 1, and we assume equal proportions. The parameter $\beta_{E}$ has no direct consequence on the inference and was set to 0.3 throughout. The error term was simulated from either a standard normal distribution (i.e. with mean 0 and variance 1), a student’s *t*-distribution with degrees of freedom 5, or an exponential distribution with rate parameter 1.

The simulation was repeated 10,000 times by sampling from the error distribution each time, while keeping the linear term in Equation (1) and the genotypes fixed. The simulation was performed on an iMac machine with 3.2GHz Quad-Core Intel Core i5 and 8GB of 1600 MHz DDR3 memory.

**Results**

The sex-stratified mean and variance conditions give a total of 9 combinations of simulated quantitative trait (Figure 1-3). As expected, for *gL* using the default 3-degrees of freedom test (Chen et al., 2021), the empirical type I error rates under all scenarios are well-controlled (Table 1), with the exception of errors following a student’s *t* and an exponential distribution when the MAF was 0.1. Meanwhile, the proposed scale test *gS* (Deng et al., 2019) had excellent empirical type I error control. The interesting phenomenon occurs when we computed the gJLS *p*-value via Fisher’s method that combines the location and scale tests, which are separately well controlled.

**Table 1.** Empirical type I error rate of the location, scale and the joint location scale *p*-values under different simulation designs.

| **Distribution** | **MAF** | **gL** | **gS** | **gJLS** | $\boldsymbol{\rho}$^1^ | **Correlation *p*-value**^2^ |
| --- | --- | --- | --- | --- | --- | --- |
| normal | 0.1 | 0.0493 | 0.0451 | 0.0482 | 0.00 | 0.658 |
|  | 0.2 | 0.049 | 0.0474 | 0.0488 | 0.00 | 0.957 |
|  | 0.3 | 0.0469 | 0.0486 | 0.0439 | -0.01 | 0.558 |
|  | 0.4 | 0.0457 | 0.0478 | 0.0455 | -0.01 | 0.590 |
|  | 0.45 | 0.0473 | 0.0472 | 0.0457 | -0.01 | 0.535 |
| student’s *t*  (5 df) | 0.1 | 0.0536 | 0.0487 | 0.0529 | 0.00 | 0.649 |
|  | 0.2 | 0.0489 | 0.0476 | 0.046 | 0.01 | 0.326 |
|  | 0.3 | 0.0489 | 0.0466 | 0.0465 | 0.00 | 0.650 |
|  | 0.4 | 0.0557 | 0.0501 | 0.054 | 0.03 | 0.004 |
|  | 0.45 | 0.0476 | 0.0451 | 0.0483 | 0.00 | 0.976 |
| exponential | 0.1 | 0.0567 | 0.0425 | 0.0737 | 0.50 | 0.000 |
|  | 0.2 | 0.0453 | 0.0467 | 0.0707 | 0.52 | 0.000 |
|  | 0.3 | 0.0492 | 0.0488 | 0.0785 | 0.52 | 0.000 |
|  | 0.4 | 0.0482 | 0.0454 | 0.0749 | 0.52 | 0.000 |
|  | 0.45 | 0.0461 | 0.0472 | 0.0745 | 0.51 | 0.000 |

^1^The correlation between location and scale *p*-values is captured by a rank-based correlation coefficient (Spearman’s rho) and **^2^** the associated *p*-value is reported under the null hypothesis that the true rho is equal to zero.

Perhaps unsurprisingly, when the error term is skewed (e.g. following an exponential distribution), the combined gJLS *p*-value is persistently inflated (Table 1). This can be explained by the fact that the location and scale *p*-values were correlated as measured by Spearman’s rho ($\boldsymbol{\rho}$), a rank-based correlation coefficient. Meanwhile, though the student’s *t* distribution presents a departure from normality, the fat-tail had minimal impact on the gJLS *p*-value. Results from this simulation suggest that the gJLS can be reliably applied to quantitative traits that are at least symmetrically distributed where the location and scale test are separately well controlled in terms of type I error rate. The MAF of the SNP also had some influence on both the location and scale test results, which makes the combined gJLS *p*-values also sensitive to small MAFs as observed here and previously in Soave et al., (2015, 2017).

References

Chen, B., Craiu, R. V., Strug, L. J., & Sun, L. (2021). The X factor: A robust and powerful approach to X-chromosome-inclusive whole-genome association studies. *Genetic epidemiology*, **10**.1002/gepi.22422. Advance online publication. <https://doi.org/10.1002/gepi.22422>

Deng, W. Q., Mao, S., Kalnapenkis, A., Esko, T., Mägi, R., Paré, G., & Sun, L. (2019). Analytical strategies to include the X-chromosome in variance heterogeneity analyses: Evidence for trait-specific polygenic variance structure. *Genetic epidemiology*, ***43***(7), 815–830. <https://doi.org/10.1002/gepi.22247>.

Lindström S, Yen YC, Spiegelman D, Kraft P. (2009) The impact of gene-environment dependence and misclassification in genetic association studies incorporating gene-environment interactions. *Human Heredity*. **68**(3):171-181. <https://doi.org/10.1159/000224637>

Roslin, N. M., Weili, L., Paterson, A. D. & Strug, L. J. (2016). Quality control analysis of the 1000 Genomes Project Omni2.5 genotypes. *BioRxiv*, <https://doi.org/10.1101/078600>.

Soave, D., Corvol, H., Panjwani, N., Gong, J., Li, W., Boëlle, P. Y., Durie, P. R., Paterson, A. D., Rommens, J. M., Strug, L. J., & Sun, L. (2015). A Joint Location-Scale Test Improves Power to Detect Associated SNPs, Gene Sets, and Pathways. *American journal of human genetics*.  ***97***(1), 125–138. <https://doi.org/10.1016/j.ajhg.2015.05.015>.

Soave D, Sun L. (2017). A generalized Levene's scale test for variance heterogeneity in the presence of sample correlation and group uncertainty. *Biometrics*. **73**(3):960-971.

<https://doi.org/10.1111/biom.12651>.

**Fig. 1. Distribution of simulated quantitative traits with normal error.** The male- and female-specific densities are shown in red and blue, respectively. The headers indicate the model parameters values used for each simulated scenario.

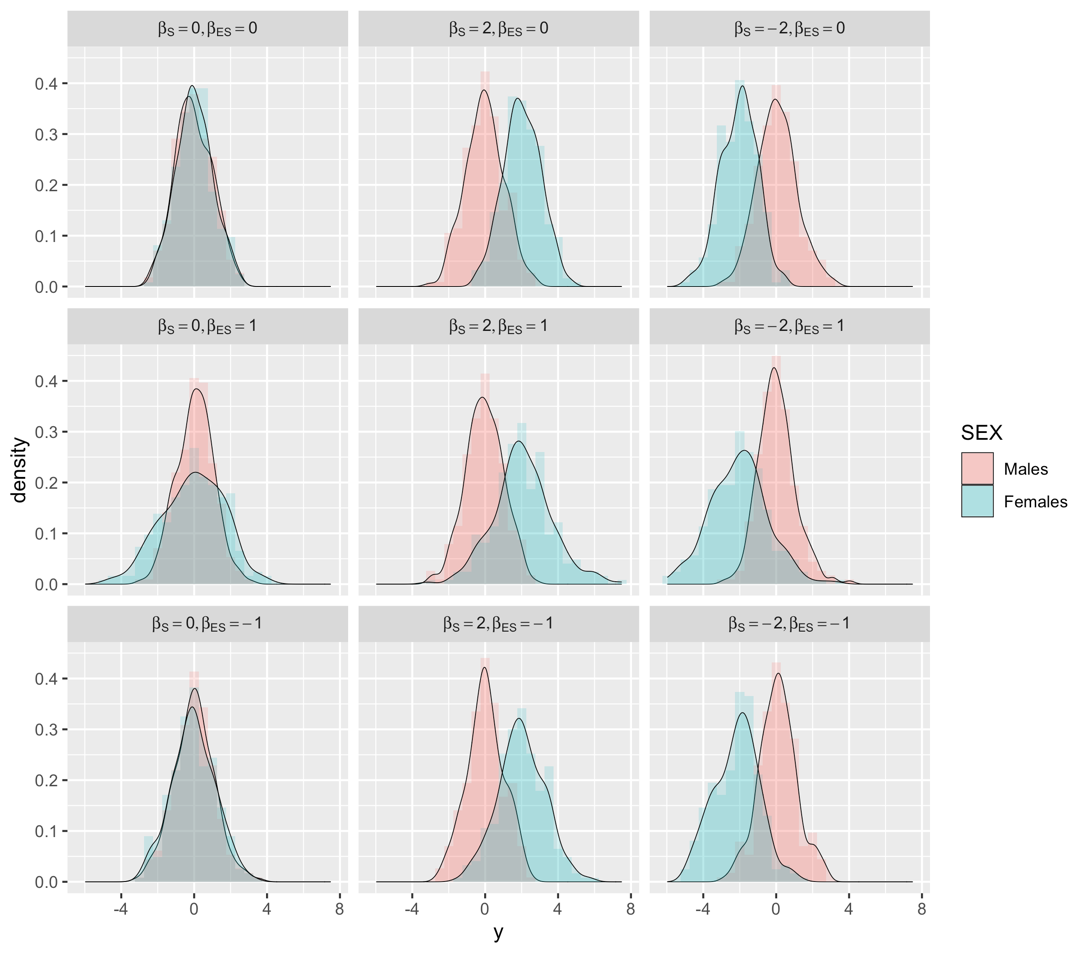

**Fig. 2. Distribution of simulated quantitative traits with t-distributed error.** The male- and female-specific densities are shown in red and blue, respectively. The headers indicate the model parameters values used for each simulated scenario

.

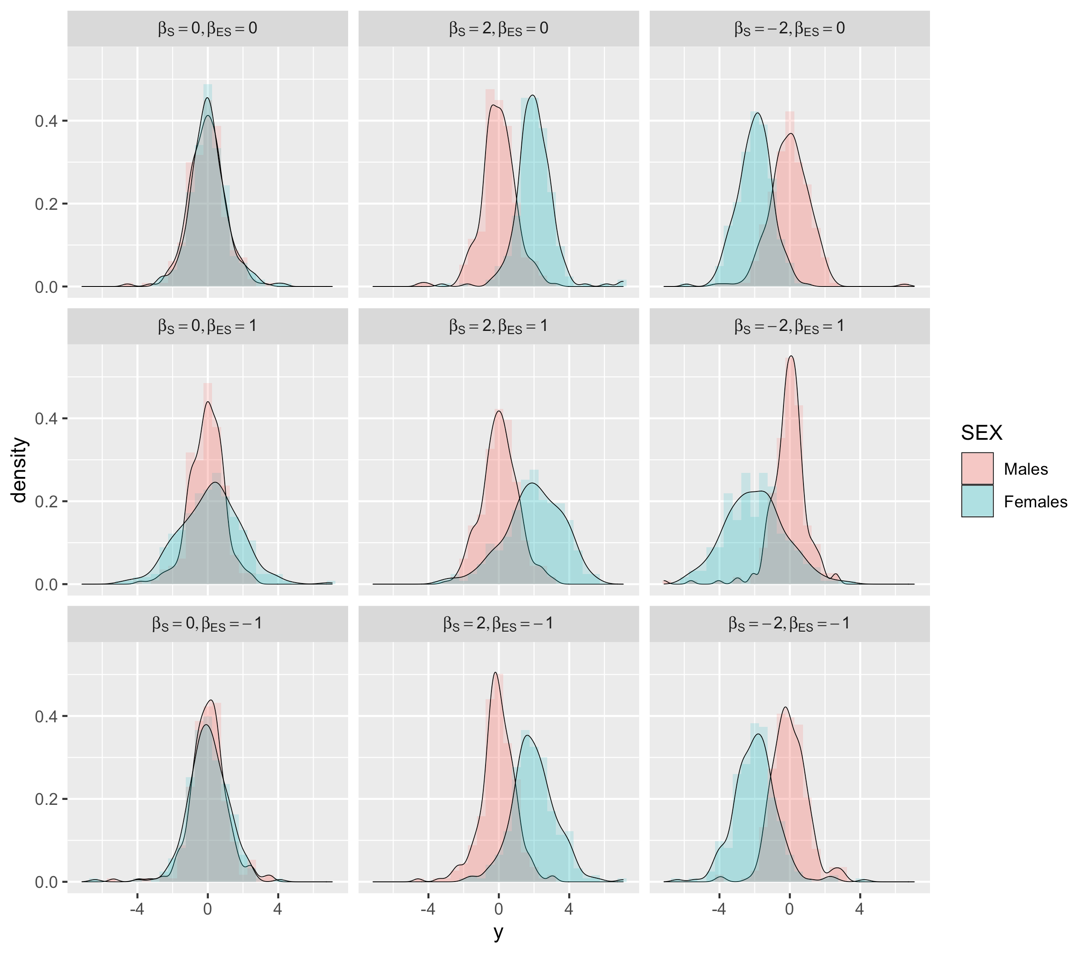

**Fig. 3. Distribution of simulated quantitative traits with an exponentially distributed error.** The male- and female-specific densities are shown in red and blue, respectively. The headers indicate the model parameters values used for each simulated scenario.

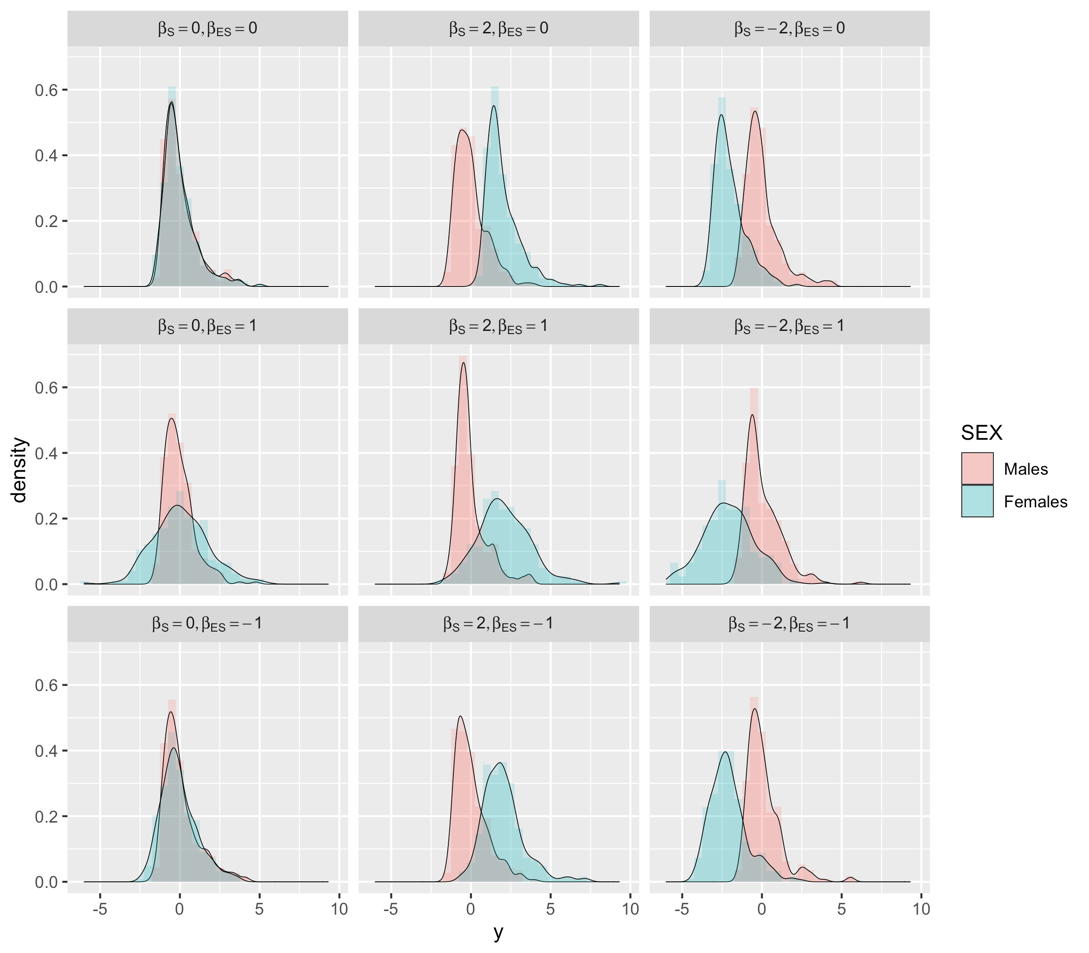
