## Supplementary figures and images for "gJLS2: An R package for generalized joint location and scale analysis in X-inclusive genome-wide association studies"

### Supplemental Figure 1

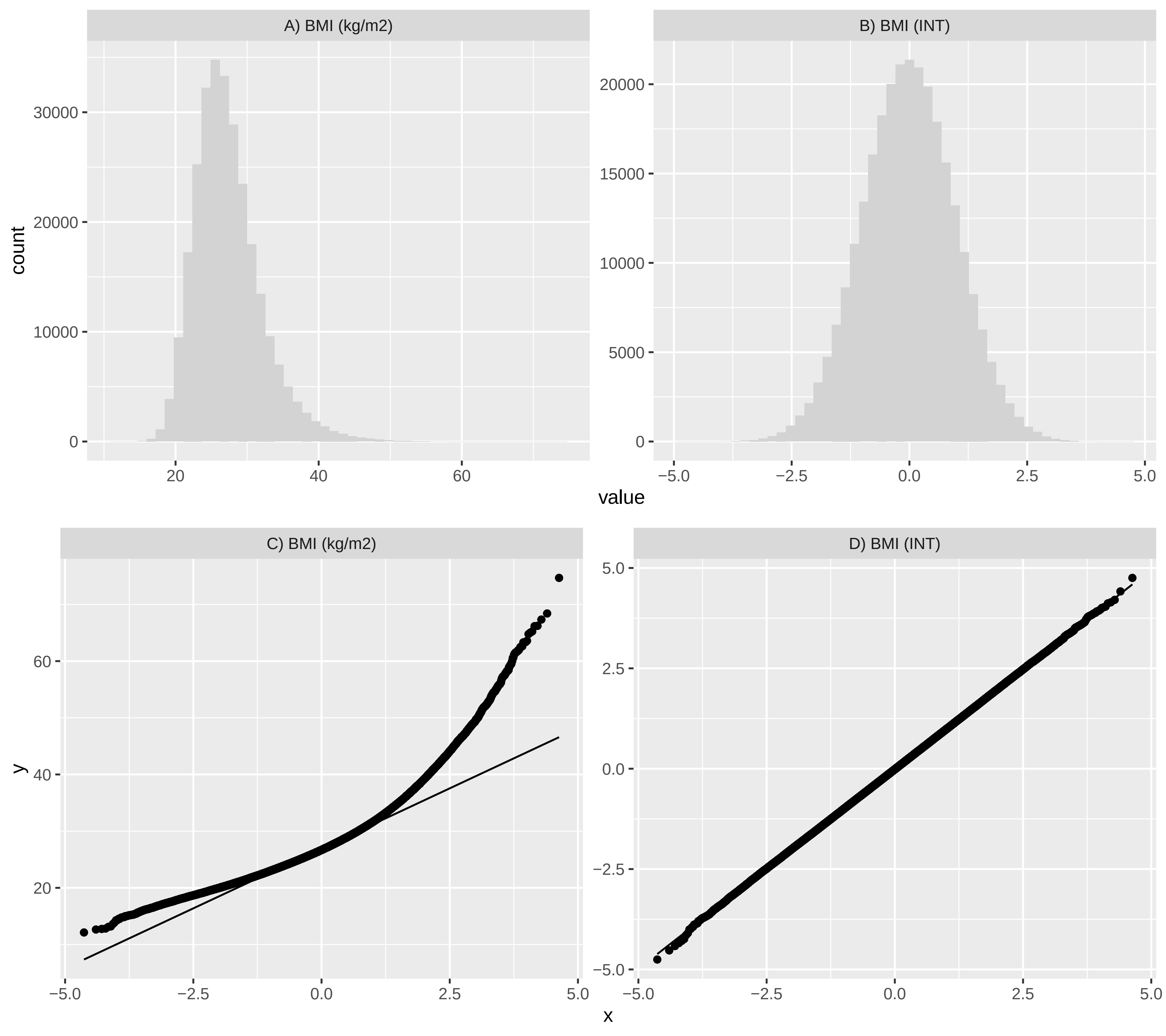

### Supplemental Figure 2

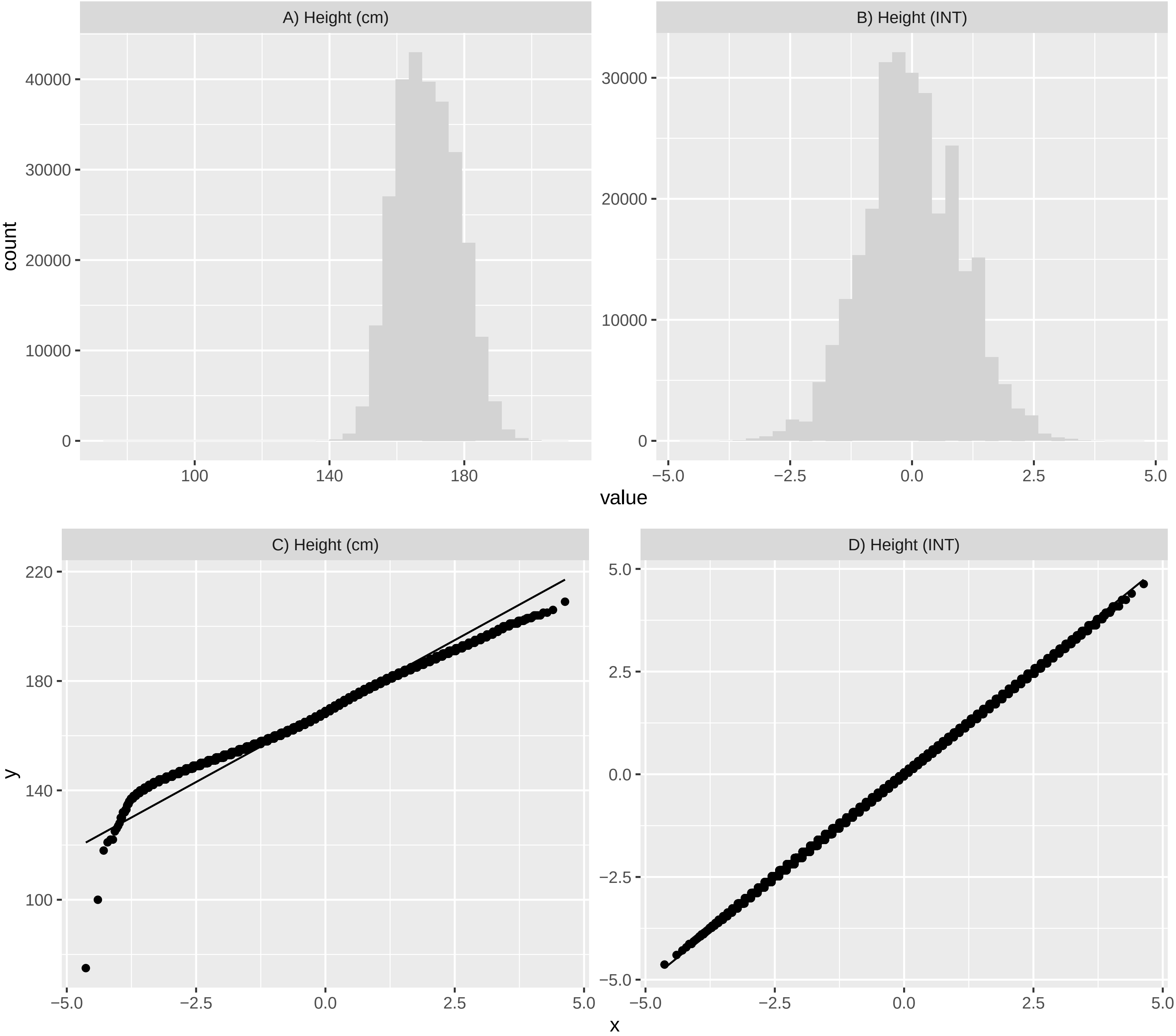

### Supplemental Figure 3

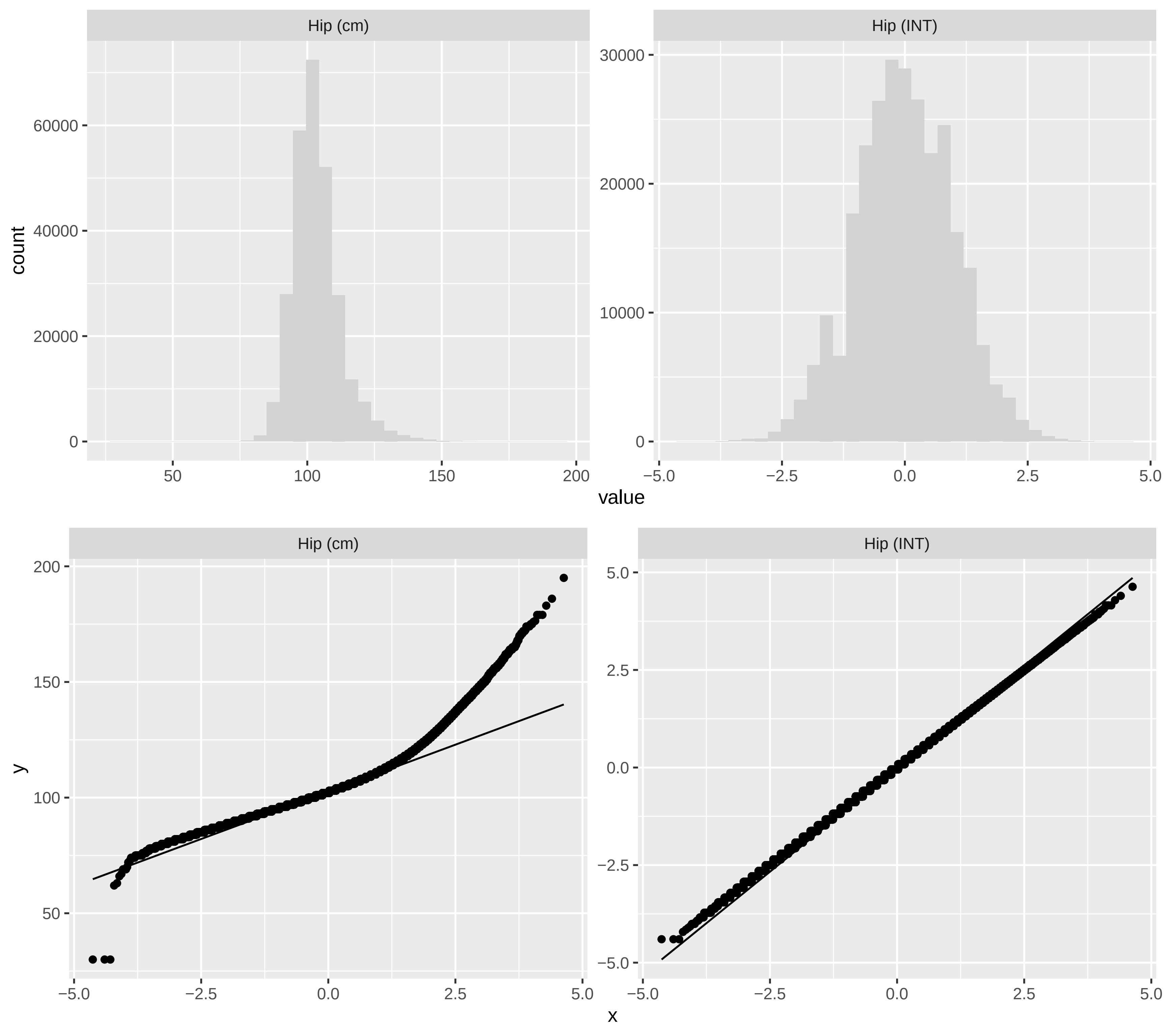

### Supplemental Figure 4

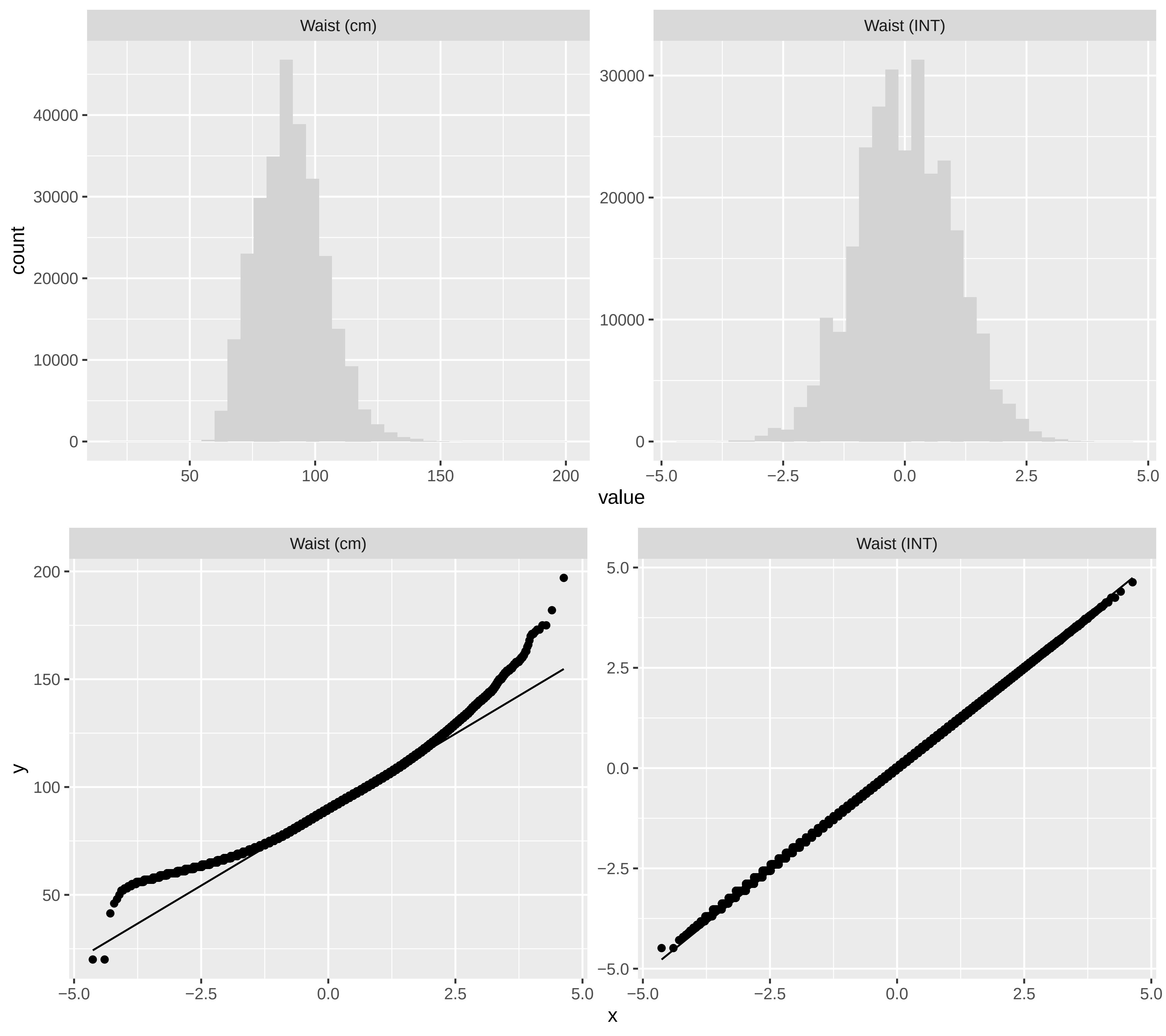

### Supplemental Figure 5

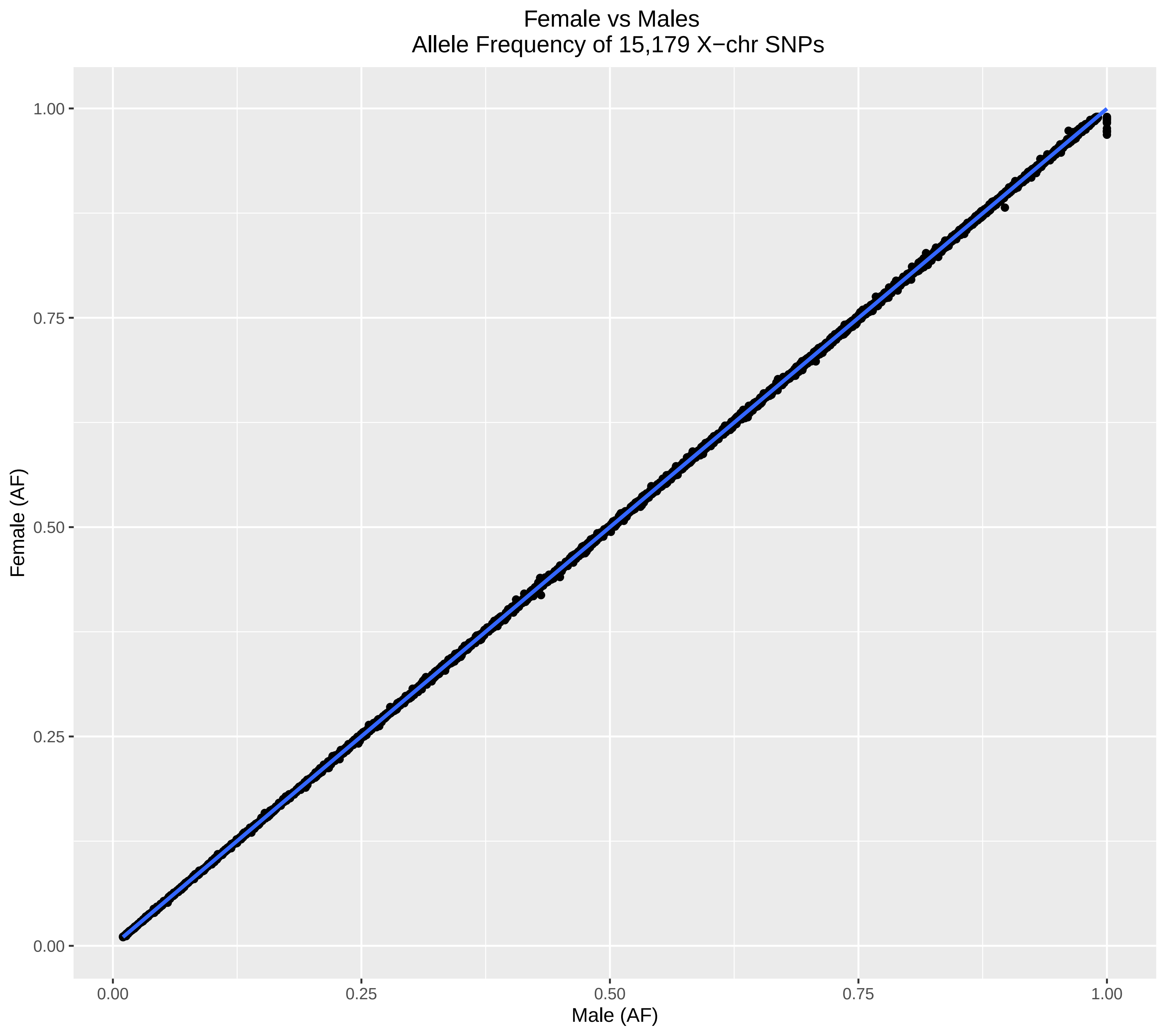

### Supplemental Figure 6

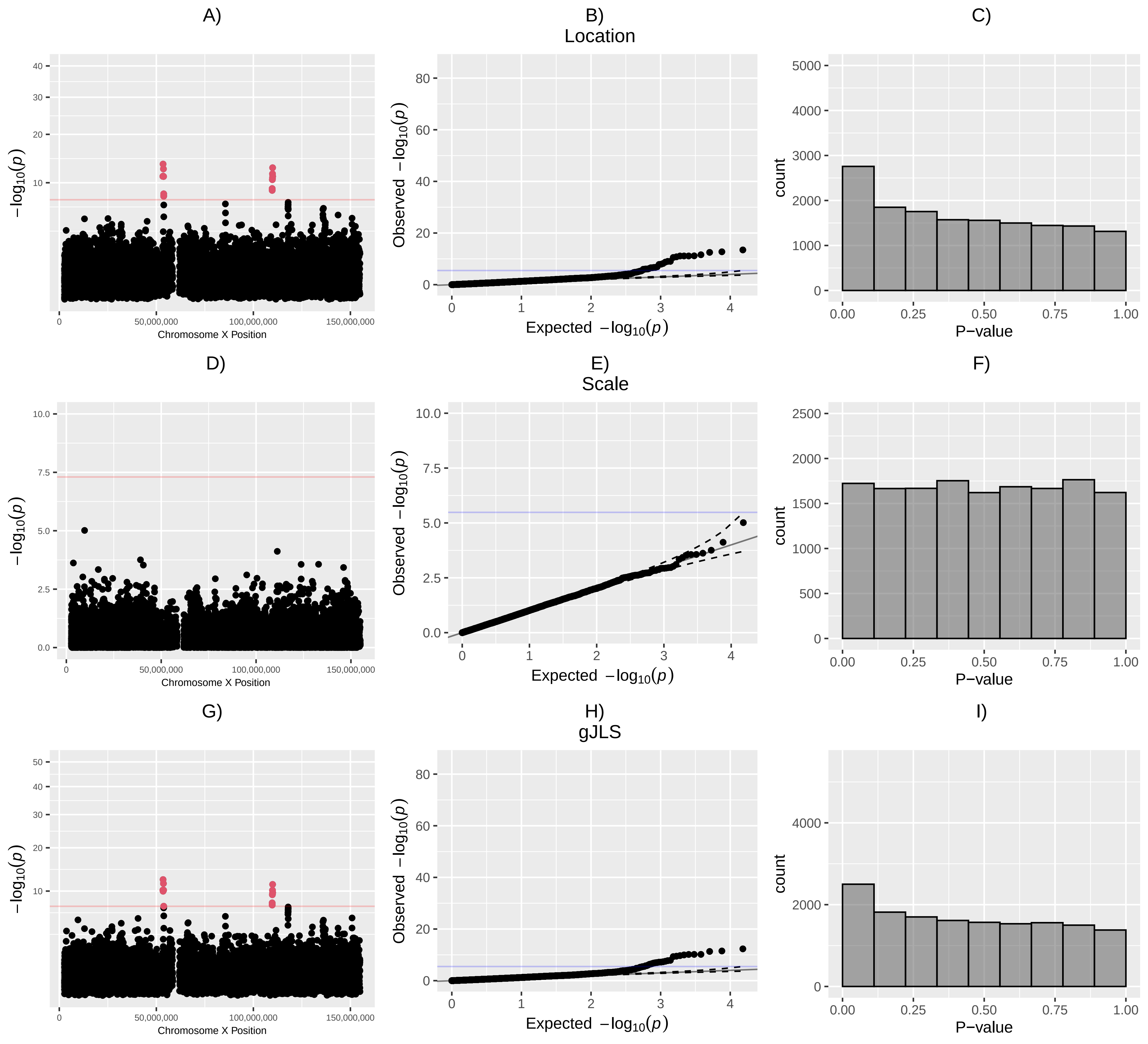

### Supplemental Figure 7

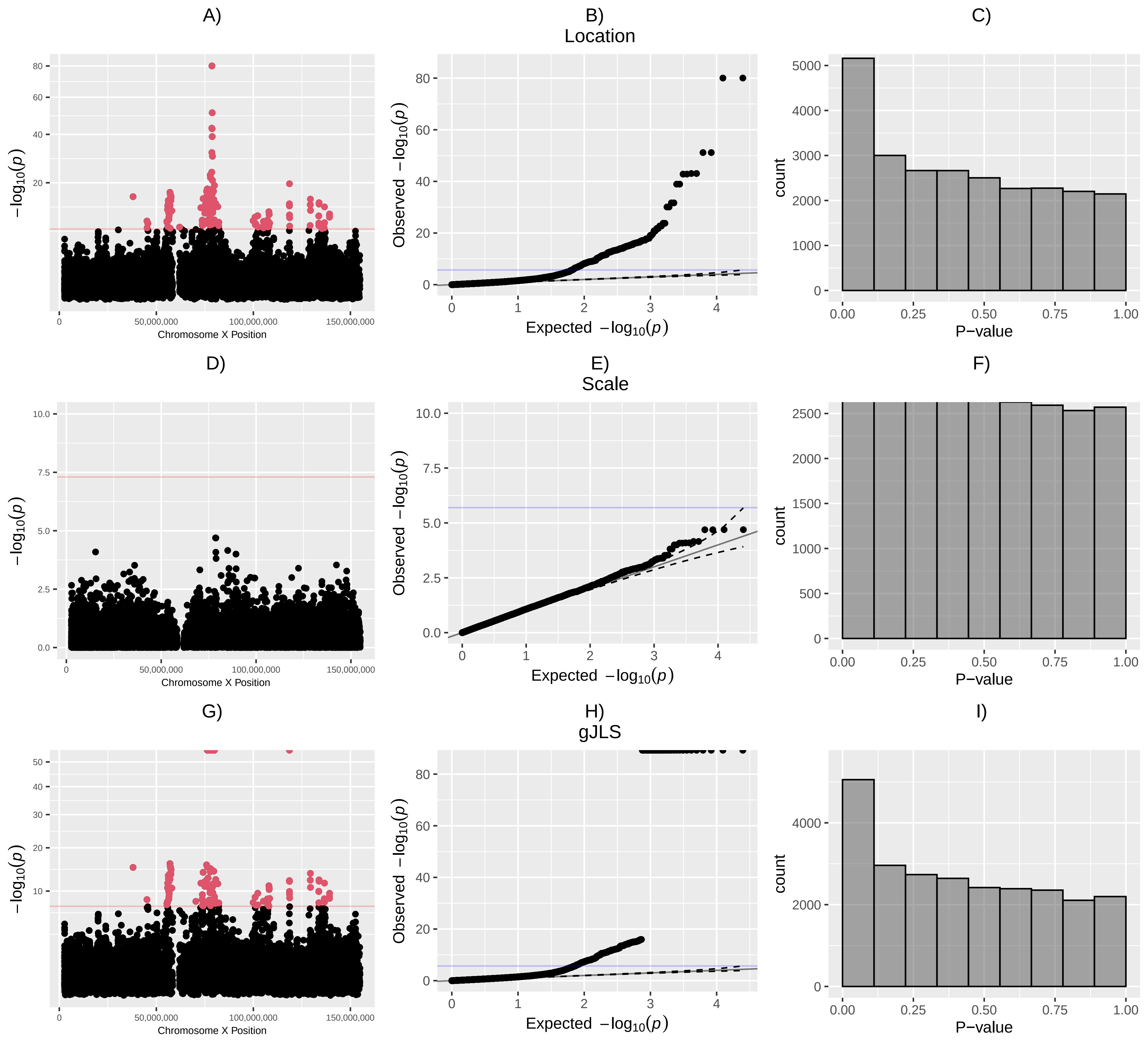

### Supplemental Figure 8

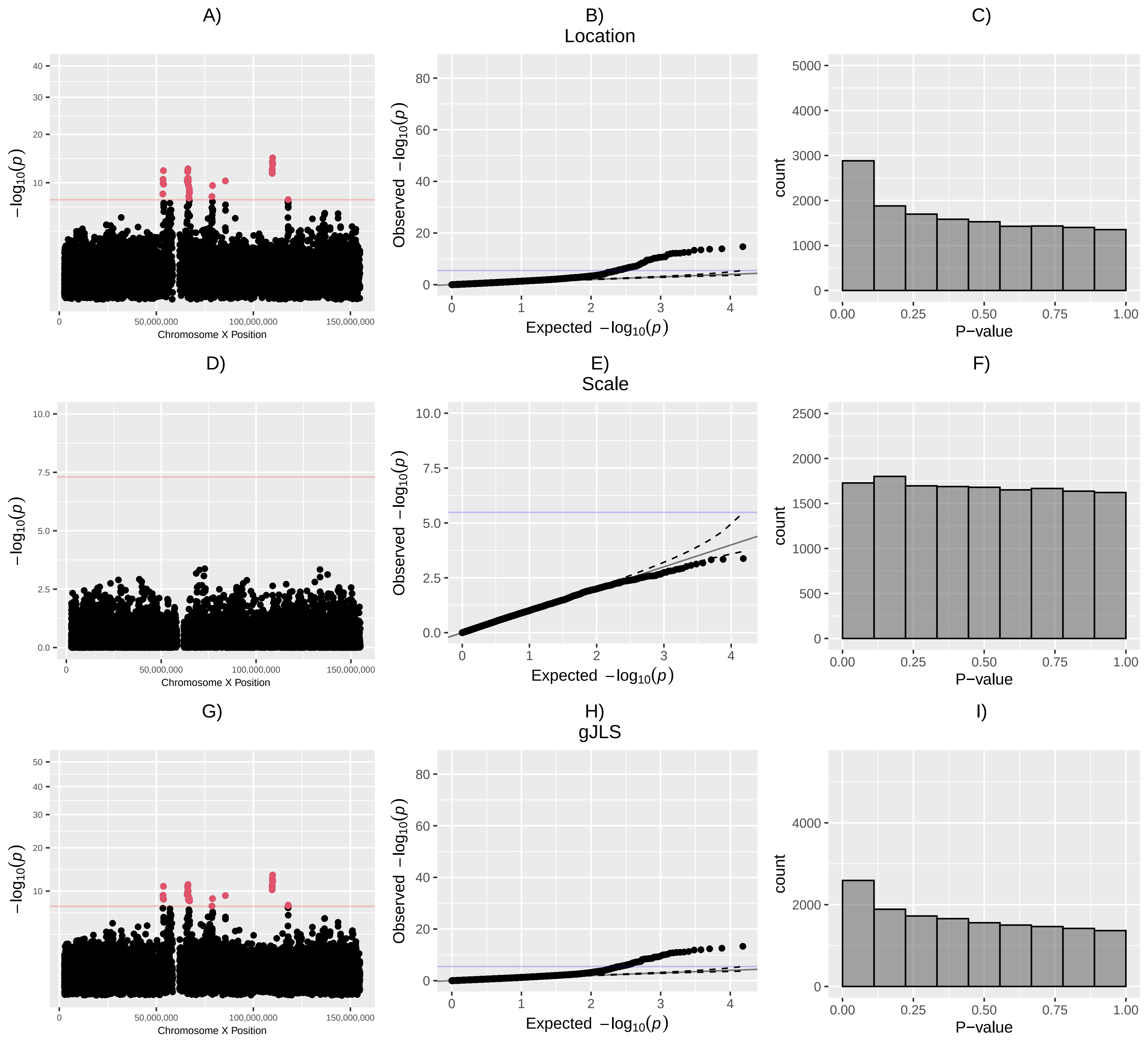

### Supplemental Figure 10

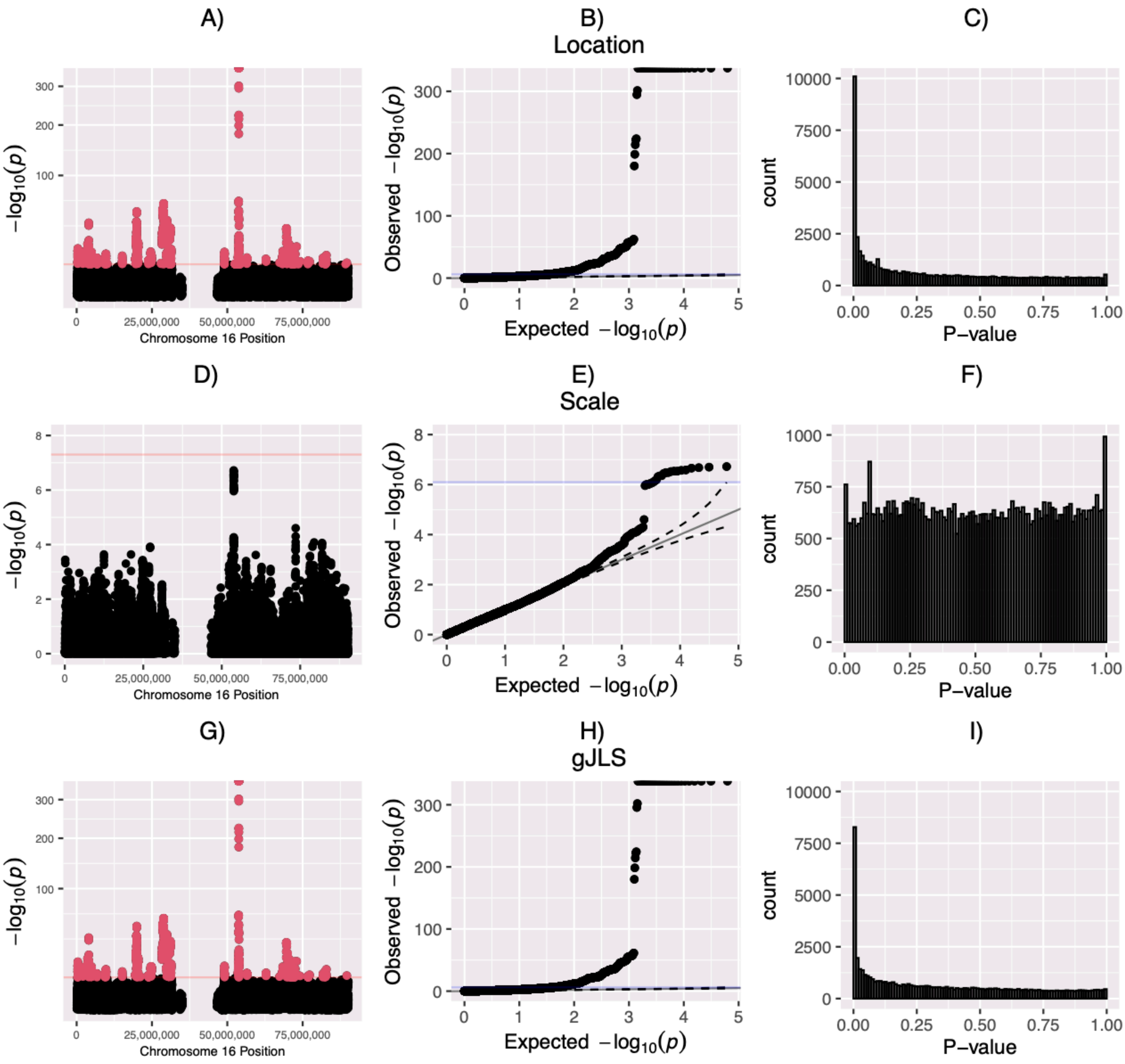

### Supplemental Figure 11

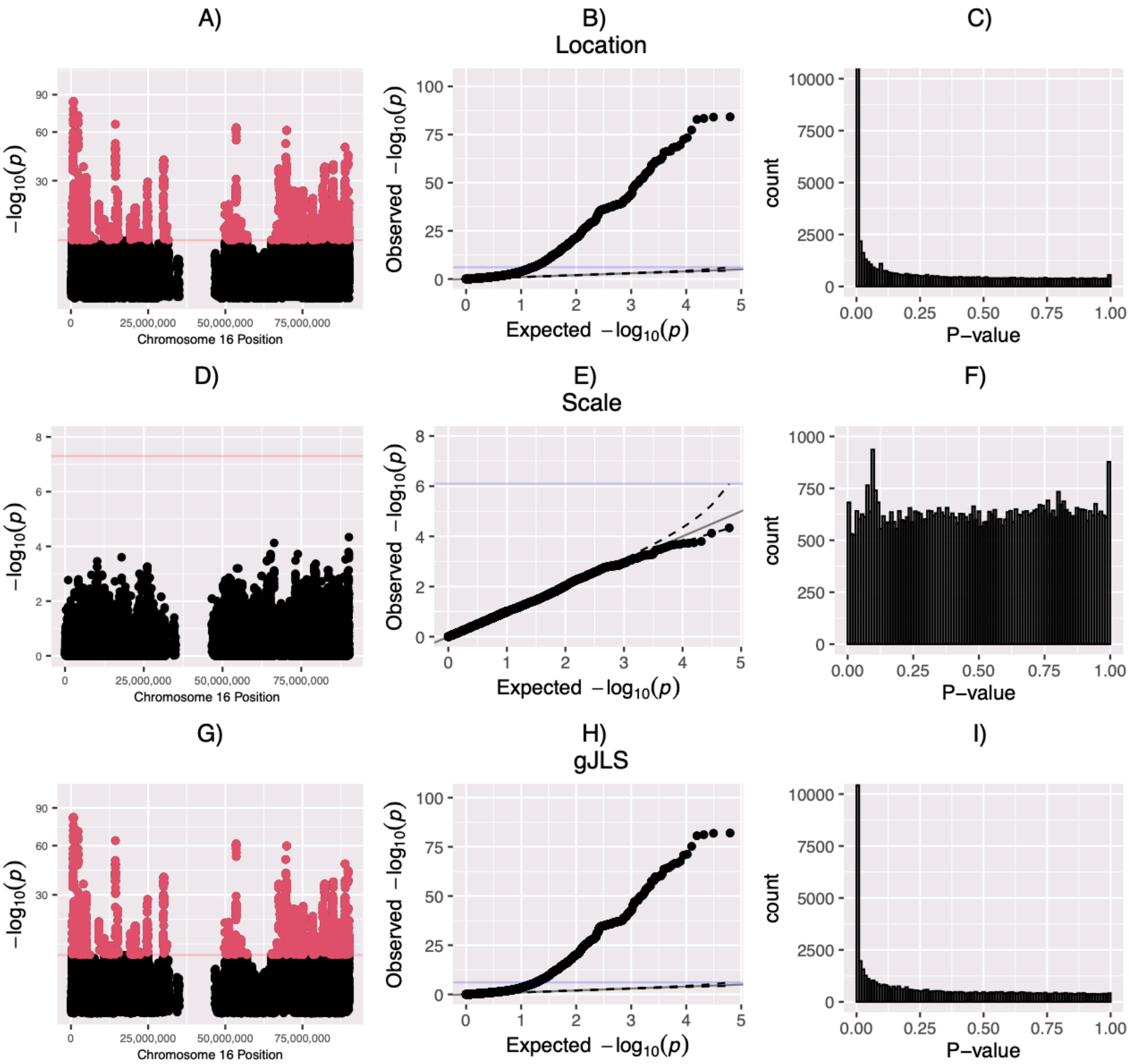
